# The Structural Basis for Glycerol Permeation by human AQP7

**DOI:** 10.1101/858332

**Authors:** Li Zhang, Deqiang Yao, Fu Zhou, Qing Zhang, Ying Xia, Qian Wang, An Qin, Jie Zhao, Dianfan Li, Lu Zhou, Yu Cao

**Affiliations:** CAS Center for Excellence on Molecular Cell Science, Institute of Biochemistry and Cell Biology, Chinese Academy of Sciences; University of Chinese Academy of Sciences, 333 Haike Road, Shanghai 201210, China; Institute of Precision Medicine, the Ninth People’s Hospital, Shanghai Jiao Tong University School of Medicine, 115 Jinzun Road, Shanghai 200125, China; iHuman Institute, ShanghaiTech University, Shanghai 201210, China (present affiliation for D.Y.); Department of Orthopaedics, Shanghai Key Laboratory of Orthopaedic Implant, Shanghai Ninth People’s Hospital, Shanghai Jiao Tong University School of Medicine, Shanghai 200011, China; Department of Medicinal Chemistry, School of Pharmacy, Fudan University

**Keywords:** Glycerol Metabolism, aquaporin, membrane protein, transportation across membrane

## Abstract

Human glycerol channel AQP7 conducts glycerol release from adipocyte and entry into the cells in pancreatic islets, muscles and kidney tubule, and thus regulate glycerol metabolism in those tissues. Compared with other human aquaglyceroporins, AQP7 shows a less conserved “NPA” motif in the center cavity, and a pair of aromatic residues at Ar/R selectivity filter. To understand the structural basis for the glycerol conductance, we crystallized the human AQP7 and determined the structure at 3.7 Å. A substrate binding pocket was found near to the Ar/R filter and the bound glycerol molecule stabilized by R229. *In vivo* functional assay on human AQP7 as well as AQP3 and AQP10 demonstrated strong glycerol transportation activities at physiological condition. The human AQP7 structure reveals a fully closed conformation with its permeation pathway strictly confined by Ar/R filter at the exoplasmic side and the gate at the cytoplasmic side, and the dislocation of the residues at narrowest parts of glycerol pathway in AQP7 play a critical role in controlling the glycerol flux.

## Introduction

Glycerol, one of the most commonly found polyols in life, is involved in both lipid metabolism and carbohydrate metabolism as energy carrier and molecular backbone. Glycerol is stored in form of triacylglycerols (TAGs) in adipocytes and could be mobilized into free glycerol by lipoprotein lipase upon starvation [1]. The glycerol generated in cytosol is released into circulation and re-absorbed by liver, where it is phosphorylated into glycerol-3-phosphate (G3P) by glycerol kinase and enters the lipid synthesis or glycolytic pathways [2]. In addition to glycerol efflux, AQP7 plays a key role in glycerol uptake in kidney, muscles, pancreatic islets and male reproductive system [3-6]. Glycerol can be absorbed by muscle cells via AQP3 and 7, and utilized as energy mass via either oxidation by glycerol dehydrogenase into glyceraldehyde or phosphorylation into G3P by glycerol kinase [7, 8]. Glycerol is a small water-soluble molecule with poor permeability across lipid membranes, and thus need carriers across cell membrane to exit or enter the cell. In bacteria, *e. g., E. coli*, the glycerol is permeated by the transporter GlpT and the water channel AqpZ [9]. In eukaryotic cells, the aquaporin family conducts most of glycerol transportation across membrane. Aquaporin (AQP), or aquaglyceroporin, is a set of water/glycerol channel protein comprising of thirteen members in human genome, Aqp0-Aqp12, four of which are associated with glycerol transportation in cells: AQP3, AQP7, AQP9 and AQP10 [10]. AQP7 is highly expressed in adipose tissue, renderingthe membrane permeable to water and glycerol [11]. Adipose cells store excess nutrition in the form of TAGs and hydrolyze them into free fatty acid (FFA) and glycerol upon starvation, representing an evolutional strategy to survive fluctuation of food supply. The rapid accumulation of glycerol needs efficient cleanup to prevent cell damage caused by elevated osmatic pressure. As well, a tight regulation on transportation across membrane is required to avoid unnecessary loss of energy molecules. The functional deficiency of AQP7 results in glycerol accumulation in adipose tissues and causes obesity in adulthood in mice model with AQP7 knocked out [12, 13]. In human studies, a loss-of-function mutation, G264V, is responsible for several healthy issues such as hyperglyceroluria, psychomotor retardation and abnormal platelet [14]. Gly264 is predicted to be located in the sixth transmembrane helix of AQP7 [15] Other consequences of AQP7 deficiency includes impairment of β-cell function and insulin secretion [3, 16], as well as high risk of ventricular hypertrophy in AQP7-KO mice [5].

Although the sequence diversity among AQP family is quite high and the sequence identity among human AQP proteins ranges from about 17% to about 41%, the published water-AQP structures show a highly conserved folding pattern including an overall architecture of AQP homo-tetramer surrounding a 4-fold axis of symmetry perpendicular to cell membrane, and each protomer consisting of six transmembrane helices which establish four individual water permeation pathways in a tetramer [17-21]. A dual Asn-Pro-Ala (NPA) motif in the center region of water pathway has been suggested as an ion-repelling pore that prevents charged atoms from penetrating the water channel. A recent human AQP10 structure reveals a central channel that is slightly wider than previous reported structures of water-AQPs and proposed a pH-gating mechanism by which the channel was controlled by the protonation of H80 and its interactions with E27 at intracellular vestibule [22]. As the major glycerol channel in physiological condition, AQP7 shares high sequence identity with AQP10, especially at pH-sensor domain found in AQP10. In a functional assay that monitors the cell volume change for AQP-expressing yeast cells as a function of cellular solute concentration change, AQP7 shows a pH-gating pattern in uptake assay and displays a strong glycerol-permeation activity at physiological conditions, while AQP10 only opens to glycerol at extremely acidic conditions, implying the importance of revealing the structural basis of glycerol permeation by AQP7 in understanding lipid and glycerol metabolism [22]. To investigate the molecular mechanism underlying routine glycerol transportation in-and-out of the cells, we determine the crystal structure for human AQP7 and analyzed the regulation of the glycerol uptake conducted by AQP7 and its homolog genes AQP3 and AQP10.

## Results

### The Glycerol Permeation by Aquaglyceroporins and the overall architecture of AQP7

To investigate the glycerol permeation ability of aquaglyceroporins, AQP3, AQP7, and AQP10 were tested for their transportation of ^14^C-labelled glycerol across the cell membranes of HEK293 cells overexpressing the abovementioned channels. Interestingly, cells carrying all three aquaglyceroporins displayed strong and comparable level of glycerol uptake at physiological pH (Figure 1A), which can all be efficiently inhibited by adding non-labelled glycerol but keep unchanged upon adding urea, a potential substrates for AQP3 and AQP7 [23, 24], implying a strong substrate selectivity of the AQPs tested. This result was inconsistent with a previously reported observation of inability of AQP10 to allow glycerol permeation at neutral pH revealed by a fluorescence-based assay [22].

**Figure 1.**
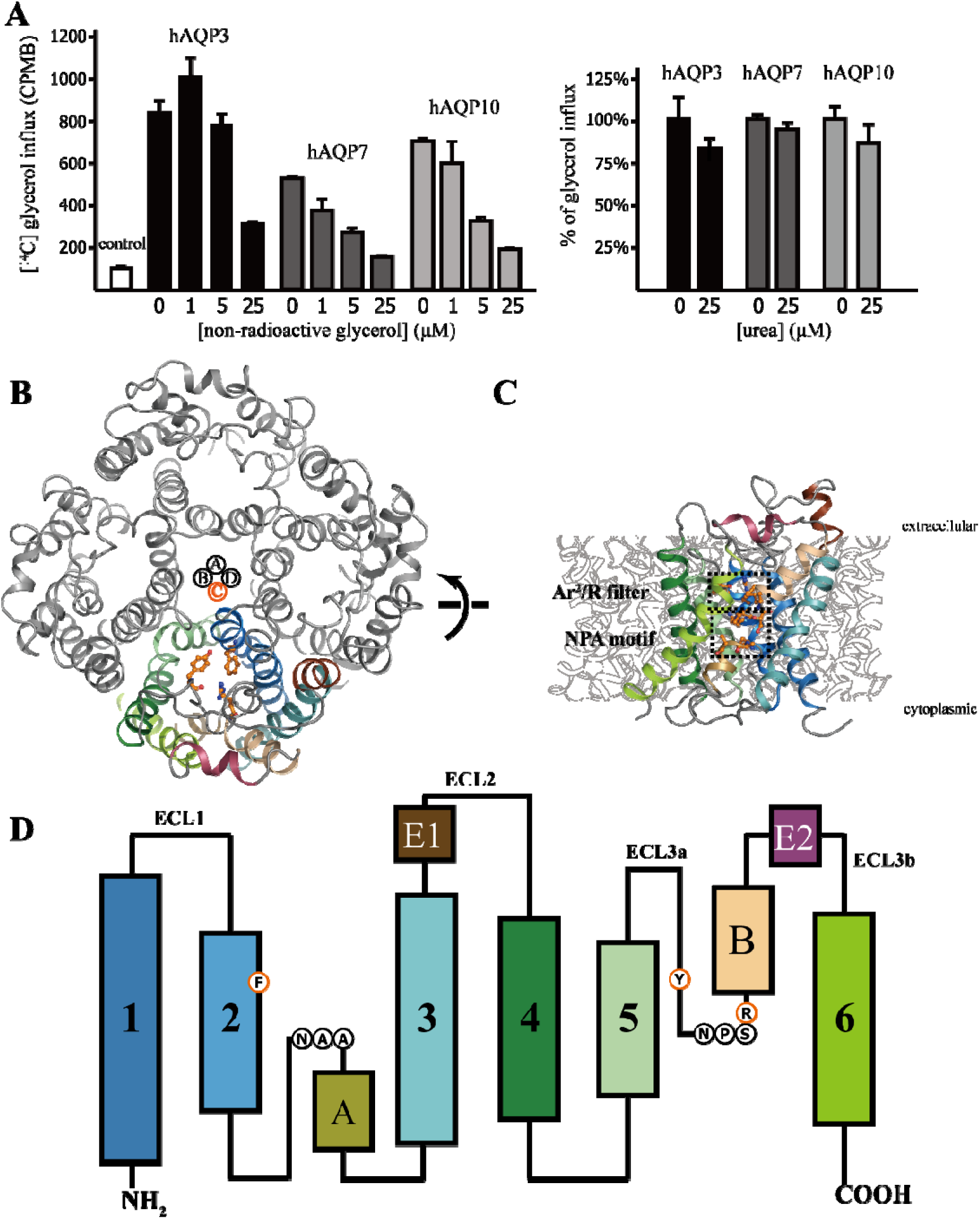
Glycerol uptake by aquaglyceroporins and the overall architecture of AQP7 tetramer. (A) Left: [^14^C]-radioactive glycerol uptake by HEK293 cells expressing human AQP3, 7 and 10 and the competitions by addition of non-radioactive glycerol at concentration indicated. The control is HEK293 cells transfected with empty vectors and represents the basal glycerol uptake. Right: The competition on glycerol uptake by 25μM of urea. (B) Tetrameric AQP7 protein viewed from the extracellular side. The four AQP7 molecules in the tetramer were designated as chain A, B, C, and D, respectively, with chain A, B and D colored in gray and C colored by transmembrane helices. The Ar^2^/R domain in chain C was shown in ball-and-stick model. (C) AQP7 viewed parallel to the membrane. The cell membrane was shown as gray block and the orientation labelled according to the calculation using PPM server (https://opm.phar.umich.edu/ppm_server). (D) Cartoon representation of AQP7 monomer topology. The TM helices 1-6, extracellular helices E1 and E2, and membrane-embed helices A and B were colored according to the same scheme as in (A) and (B). The residues in Ar^2^/R and “NPA” motifs were marked as singled-letter codes in circles. All structure graphs in this paper were produced using PyMOL (The PyMOL Molecular Graphics System, Version 1.9 Schrödinger, LLC.) and UCSF Chimera (The UCSF Resource for Biocomputing, Visualization, and Informatics, version 1.13.1) [30].

To further understand the glycerol transportation by aqp channels, human AQP7 was purified from insect cell/baculoviral expression system. The purified protein displayed an apparent molecular weight of about 120 kDa on size-exclusion gel filtration which was mostly consistent with a homo tetramer assembly. Initial crystallization trials yielded thin-plate crystals diffracting to about 10 Å resolution. Extensive efforts to improve diffraction ability of full length AQP7 crystals failed but trypsin digestion of AQP7 to remove its flexible C-terminal region successfully produced a compact protein core which yielded 3-D crystal and diffracted to 3.7 Å resolution, allowing the structure determination of AQP7 by molecular replacement using the model PDB 6F7H. As shown in Fig. 1, human AQP7 adapts a typical AQP fold. The tetrameric AQP7 proteins assemble into a square-shape transmembrane helical bundle and each protomer forms an individual glycerol channel. The homotetrameric architecture is maintained with the interactions among the TM1&2 with the TM5 from neighboring AQP7 molecule, as well as the interaction of TM1-TM6 (from neighboring AQP7 protomer) at cytoplasmic region and TM3-TM4 (from neighboring AQP7 protomer) at exoplasmic region.

Due to the 4-fold symmetry from homotetramer, most AQP proteins crystallized in space groups with high symmetry such as P4_2_ (human AQP2, PDB ID 4NEF), P42_1_2 (human AQP4, PDB ID 3GD8), and I422 (human AQP1, PDB ID 4CSK; Plasmodium falciparum AQP, PDB ID 3C02; Escherichia coli, PDB ID 1FX8). However, the four-fold symmetry was slightly impaired by the conformational variation among four AQP7 monomers, resulting in the loss of 4-fold symmetry and the diffraction limitation of APQ7 crystal. In the present AQP7 crystal structure, the electron density for chain A and C was well solved, while density for chain B and D showed relatively lower quality (Fig S3). We focused on the chain C in following structural analysis and discussion because it had the best quality of electron density map among all the four chains.

### The Selectivity Filter and gating residues

An AQP7 monomer was comprised with six transmembrane helices (TM1-6) and two half-membrane spanning helix-loop domains (domain A and B) excluding non-substrate molecules from entering the glycerol permeation pathway. Although the “NPA” motif is highly conserved among human AQP family and remains unchanged in aquaglyceroporins-3, 9 and 10, in AQP7 both domain A and B contain variant motif sequences, *i. e.*, “NAA” and “NPS” for domain A and B, respectively (Figure 1D). The whole molecule shows a pseudosymmetry between two sets of TM-TM-NPA motif -TM folding, and this structural repeat was also present in several other solute transporters, such as urea transporter UT-B (PDB ID 4ezc), glucose transporter GLUT1 (PDB ID 4PYP), glycerol-3-phosphate transporter GLPT (PDB ID 1PW4), and sphingolipid transporter homolog SPNS (PDB ID 6E8J) [25-28].

AQP channels usually form an aromatic-arginine residue pair (Ar/R filter) in their extracellular vestibule to restrict the passage of substrate molecules, where the diameter of permeation pathway is less than 2 Å in most of water-AQP channels but quite wide in previously reported glycerol channel AQP10 (>2.5 Å) [22]. In AQP7, the Ar/R filter resulted from F74-R229 interaction is reinforced by the presence of an extra aromatic residue Y223, thereby forming an Ar^2^/R filter (Figure 2A) with stronger cation-π interaction. This Ar^2^/R filter could also be found in GlpF and pfAQP where W-F-R form the filter, but not in most of other human AQPs except AQP3 (Fig S1). As calculated by the program HOLE, the permeation pathway is narrow (less than 2.0 Å) in diameter at both Ar^2^/R filter and a “gate” region at intracellular vestibule, representing the entrance and exit of transmembrane pathway, respectively (Figure 2B). Both Ar^2^/R filter and “gate” are primarily closed in the crystal structure, preventing the glycerol molecules from entering the pathway from either exoplasmic or cytoplasmic sides. The gate regions in both AQP7 and AQP10 were situated near to membrane-cytosol interface and comprised with a group of charged residues (e. g., R87, R106, and D191-Q192-E193 for AQP7), rendering the gate with potential pH sensitivity. To further probe the open mechanism for the Ar^2^/R filter, the glycerol uptake assay was conducted on AQP7 with mutation on the filter residues and the loops connecting with the filter. In figure 2D, it was shown that the uptake activities were reduced upon the mutations on both filter residue Y223 and loop residue N220, an critical residue mediating the interaction between ECL3a and ECL2, implying the role of these two loop in the Ar^2^/R filter open-close switch. At the intracellular gate, point mutation on any residue from D191-Q192-E193 will result in an activity reduction, while the mutation on R87 causes no significant consequence on glycerol uptake.

**Figure 2.**
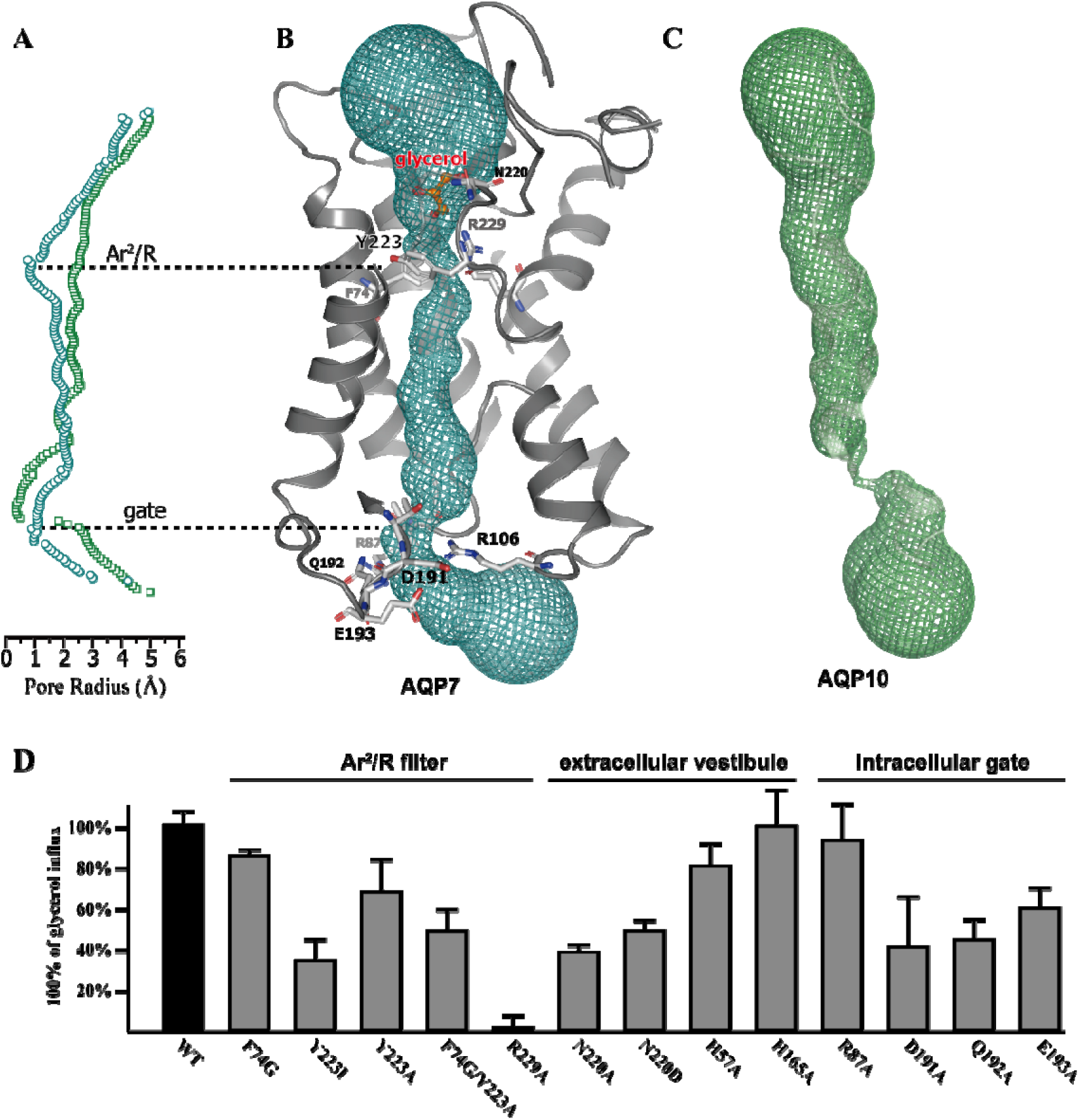
The glycerol pathway and critical residues at filter and gate. (A). the radius for inner pathway calculated by HOLE. Cyan circles showed the radius for pathway of AQP7 and green squares for AQP10. The dashed lines indicated the most narrow part and their corresponding part in glycerol pathway in B and C. (B). Glycerol permeation pathway in AQP7 as determined by HOLE was shown as cyan mesh in cartoon representation of AQP7, with the TM4 and 6 removed for a clear observation. The critical residues at Ar^2^/R and gate were shown in stick model rendered by elements. The glycerol in binding pocket above the Ar^2^/R filter was shown in ball-and-stick model and colored as red and orange. (C). AQP10 crystal structure and its glycerol permeation pathway (shown as mesh model in green). AQP10 structural model is generated using coordinates from PDB ID 6F7H. (D) The [^14^C]-radioactive glycerol uptake assay on HEK293 cell expressing AQP7 mutants. The transportation activity by mutant AQP7 was shown as percentage of activity by wildtype AQP7.

### Substrate Binding Site

Fo-Fc electron omit density map showed two blobs of density that could fit two glycerol molecules. These two sites were also autofit as glycerol by the program LigandFit in PHENIX package at the exoplasmic side of Ar^2^/R filters of chain A and C, respectively (Figure 2A and S3E) [29].The glycerol molecule is situated in a pocket built with the Ar^2^/R filter as the “bottom” and surrounded by TM1, TM2, and the extracellular loops (Figure 3A). Since no extra glycerol was added in crystallization trial, the glycerol found in crystal structure could be captured during purification where 10% w/v glycerol was applied as protein stabilizer, implying a high affinity for the glycerol to the binding pocket. As shown in figure 3B, the glycerol is caged by the carbonyl oxygens from G156, G218 and T221, and the nitrogen atom of the guanidinium group of R229. The hydrogen bond between the hydroxyl of glycerol and the guanidinium group of R229 could help the glycerol molecule stabilized at the binding pocket until the Ar^2^/R opens upon when the hydrogen bond breaks to allow the entrance of substrates. The glycerol transportation activity was significantly reduced when R229 was mutated into alanine (Figure 2D). When compared with three glycerol binding sites identified in bacterial glycerol channel GlpF and pfAQP, the glycerol binding site in AQP7 crystal structure corresponds to the G1 site with a deeper localization in the extracellular vestibule of channel, while the glycerol binding site found in AQP10 crystal structure is equivalent to the G3 site in GlpF (Figure 3C) [21, 22].

**Figure 3.**
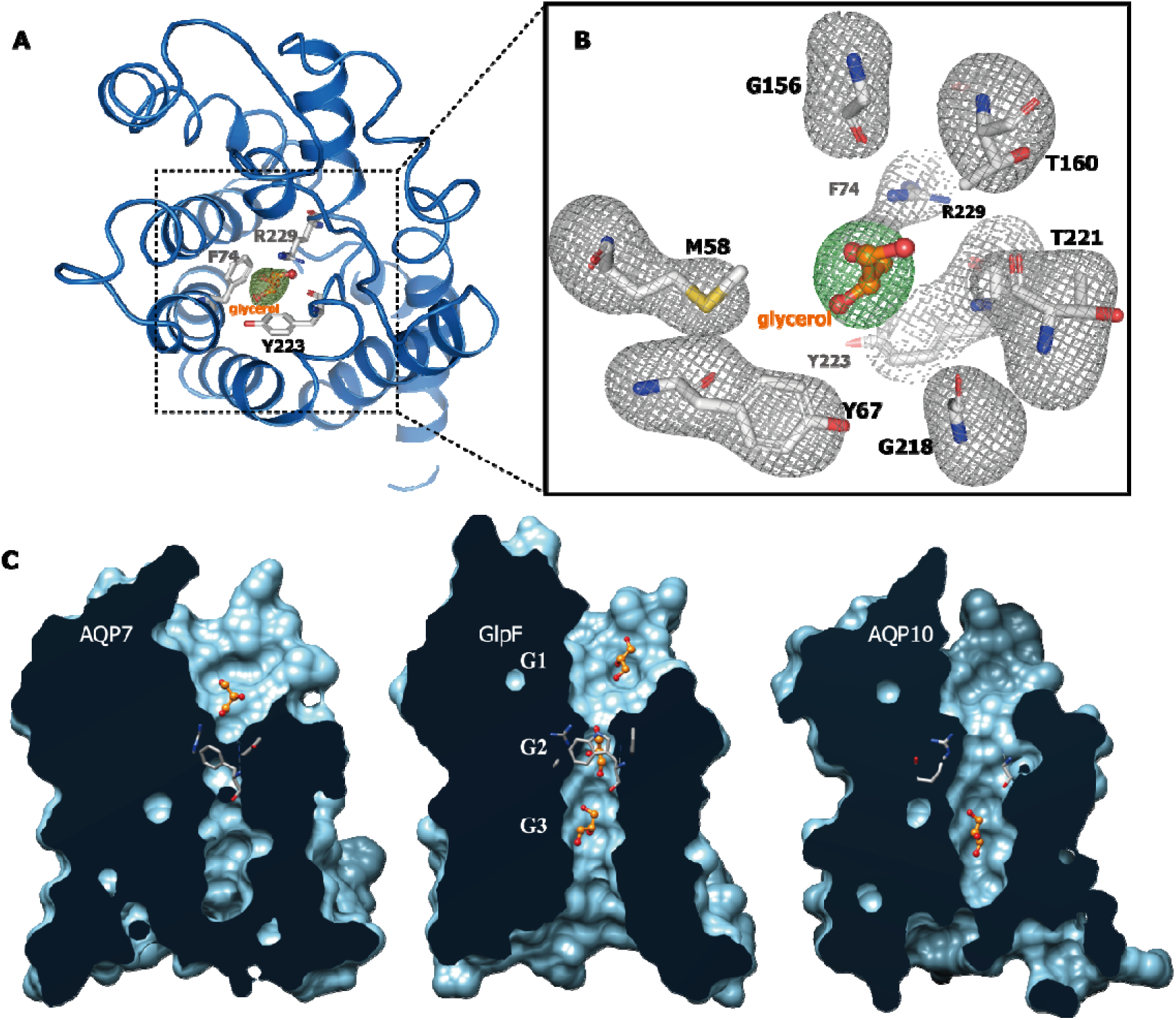
The substrate binding site in AQP7. (A). The glycerol bound and the binding site viewed from extracellular side. The glycerol was shown in ball-and-stick model and Ar^2^/R filter in stick model. (B) A close view on the binding site. The residues surrounding glycerol bound were shown in stick model. The Fo-Fc electron density was shown for glycerol (green mesh) and binding site residues (gray mesh) and contoured at 2σ. (C) A comparison among the glycerol pathway within human AQP7 (left), human AQP10 (right), and *E. coli* GlpF. The structures for all three proteins were sliced perpendicularly to membrane surface with the glycerol molecules bound shown in ball-and-stick model and Ar/R filters in stick model.

## Discussion

AQP7 is highly expressed in adipocytes and conducts glycerol efflux when intracellular glycerol level elevated by triacylglycerol metabolism. Our structure of AQP7 shows a double-shut glycerol pathway and an enhanced Ar^2^/R filter preventing the glycerol molecule in binding site from entering freely, implying a stricter regulation than that of AQP10. Indeed, among all the glycerol-AQPs in human, *Plasmodium*, and *E. coli*, AQP10 is the only aquaporin with no aromatic residues in the Ar/R gating domain, while all the others will have phenylalanine, tyrosine or tryptophan in one or two position of the Ar/R domain. In our study, the F74G mutant shows transportation activity slightly lower than wildtype, while the mutation on Y223 results in greater loss of function. The Ar^2^/R might be controlled by the ECLs and the interaction between ECL2 and ECL3 could server as switch in pathway open, probably by dragging the Y223 away from its “lock” position (Figure 4). For the gate at the intracellular end of glycerol pathway, it was shown that D191-Q192-E193 might control the substrate passage since the mutation on these two residues will cause a large drop on glycerol flux. Therefore, the function of AQP7 starts with the dislocation of Y223 from Ar^2^/R filter and D191-Q192-E193 from intracellular gate, leaving the physiological events triggering the dislocation to be further investigated.

**Figure 4.**
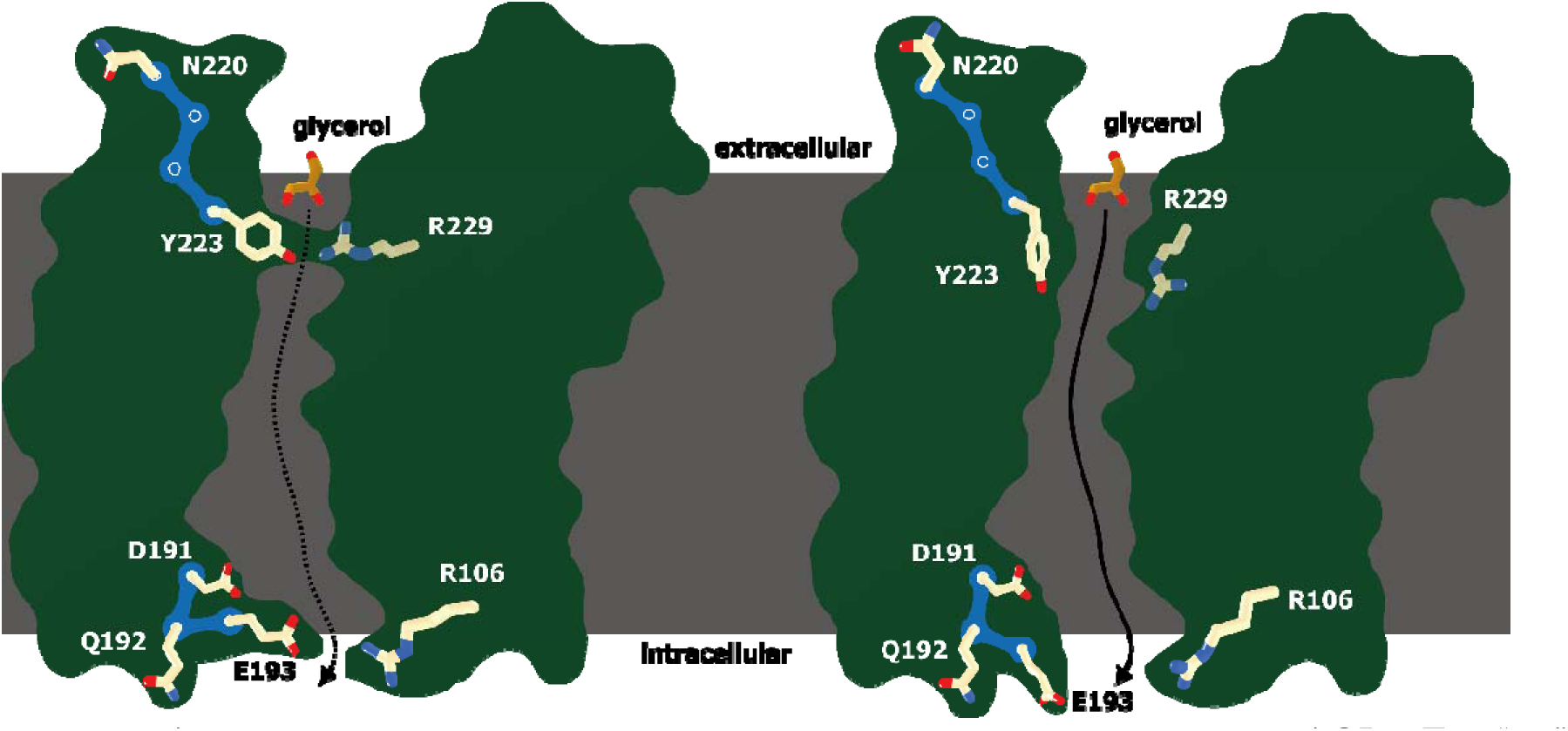
A model for the open-close mechanism of the glycerol pathway within AQP7. The “up” and “lower” gate are made with Ar2/R filter and residues Q192 and D106, respectively. The dislocation of residues N220, Q192 and D106 induce the open of glycerol pathway and thus allow the glycerol entering/leaving the cells.

AQP7 and AQP10 showed significant and opposite pH-dependence in preceding study and the glycerol transportation of AQP7 is much greater than that of AQP10 at physiological pH condition, as monitored by volume changes of yeast cell or liposome. However, in the present study, both AQP7 and 10 performed similar glycerol uptake from medium into HEK cells overexpressing AQP gene tested. We speculate the volume change of yeast cells in response to solute uptake might not be of sufficient significance in fluorescence-based assay and the rigid cell wall of the yeast might further weaken the sensitivity of measurement. Nevertheless, assuming both assay systems could properly reflect the glycerol transportation capacity of the aquaporin molecules tested, an unknown intracellular factor might exist and function in a species-specific way to facilitate AQPs, especially AQP10, to open the gate for glycerol entrance.

## Materials and Methods

### Cloning and purification of human AQP7 protein

The human AQP7 gene (Uniprot ID O14520) was codon-optimized and synthesized from GenScript, China. For expression AQP7, the full-length gene was subcloned into a modified pFastBac Dual (Invitrogen) for expression in baculovirus-insect cells with a C-terminal HRV 3C protease recognition site followed by a 2× Strep tag. The recombinant baculovirus encoding AQP7 gene was generated and infected into Spodoptera frugiperda cell line Sf9 for overexpression of AQ7 protein. The Sf9 cells were harvested 48-72 h after infection and collected by centrifugation (1500 g, 15 min, 25°C). To lyse the cells, the pellets were re-suspended with ice cooled buffer containing 10 mM NaCl, 10 mM HEPES (pH 7.5), 0.1 mg/ml DNase Ⅰ, 2 mM MgCl_2_ and 1 mM phenylmethylsulfonyl fluoride (PMSF), and the cell debris was harvested by centrifugation (45,000 g, 25 min, 4°C). To further isolate the cell membranes, the pellets were re-suspended with ice cooled buffer containing 1 M NaCl, 25 mM HEPES, pH 7.5, 0.2 mg/ml DNase Ⅰ, 2 mM MgCl_2_ and 1 mM PMSF, and homogenized with dounce homogenizer. The membranes debris were collected by centrifugation (45,000 g, 30 min, 4°C) and solubilized in lysis buffer containing 150 mM NaCl, 20 mM HEPES (pH 7.5), 0.1 mg/ml DNase Ⅰ, 2 mM MgCl_2_, 2 mg/ml iodoacetamide, and 30 mM n-Dodecyl-β-D-Maltopyranoside (DDM), supplemented with EDTA-free protease inhibitor cocktail (MedChemExpress). The solubilization slurry was incubated at 4°C with gentle stir and clarified by centrifugation (45,000 g, 45 min, 4°C). The AQP7 protein was then purified by affinity chromatography using Strep-Tactin Superflow (IBA Lifesciences). The AQP7 was further purified with gel-filtration using superdex 200 10/300 Gel column (GE lifescience) pre-equalized in a buffer containing 150 mM NaCl, 20 mM HEPES (pH 7.5), 1 mM DDM. To produce AQP7 protein core, AQP7 eluted from Strep-Tactin resin was partially digested with trypsin protease at 4°C for 30 min in a AQP7/trypsin ratio of 50:1 by weight. The proteolysis reaction was terminated by adding 1 mM PMSF and the AQP7 core was then subjected to gel-filtration in the same way to full length protein. The peak fractions containing AQP7 full length or core were pooled and concentrated to 10 mg/ml for crystallization trial.

### Crystallization

Both vapor diffusion method and liquid cubic phase method were tested for crystallization and the crystals of both full length and trypsin digested AQP7 were grown by vapor diffusion at 4□. and harvested in 3-5 days by directly flash freezing into liquid nitrogen. The crystals with diffraction capacity were grown by mixing trypsin digested AQP7 with a solution containing 30% v/v PEG400, 50 mM CAPS, pH 10.0, 0.16% w/v 1,5-Naphthalenedisulfonic acid disodium, 0.16% w/v Naphthalene-1,3,6-trisulfonic acid trisodium, and 0.16% 1,4-piperazinediethanesulfonic acid in a ratio of 1:1. The crystals were flash frozen into liquid nitrogen after cryo-protected in protectant drops. The protectant drops were produced by mixing 1 μl trypsin digested AQP7 with 1 μl high-PEG solution containing 34.6% v/v PEG400 and 50 mM CAPS, pH 10.0 and pre-equalized with high PEG solution in same way as vapor diffusion crystallization trial.

### Data collection and structure determination

Diffraction data were collected at the Shanghai Synchrotron Radiation Facility (SSRF) beamlines BL18U1 and BL19U1. The data were collected and processed with HKL2000. The crystallographic parameters and data collection statistics are given in Table S1. AQP10 structure (PDB ID 6F7H) was used as search model and the molecular replacement and initial model building were performed in Phenix. Iterative cycles of refinement were carried out using PHENIX, Refmac5 and Coot.

### *in vivo* glycerol uptake assay

Human Embryonic Kidney 293T cell (HEK293T) adherent culture overexpressing AQP genes was used to measure the transportation of ^14^C-radio labelled glycerol molecules (^14^C-glycerol). Human AQP genes (AQP3, AQP7 and AQP10) were subcloned into a transient overexpression vector modified from pcDNA3.4 and transfected into HEK293T cells in 6-well plates. The transfected

HEK293T cells were removed from 37□-incubator 36-42 h after transfection and gently washed twice with assay buffer containing 137 mM NaCl, 2.7 mM KCl, 10 mM Na_2_HPO_4_, 2 mM KH_2_PO_4_ (pH=7.4). The glycerol uptake was started by adding glycerol solution mixed from ^14^C-glycerol and unlabeled glycerol to varied final concentrations and the reaction mixtures were incubated at 37□ for designated duration until terminated by aspirating solution off and quickly washing the adherent cells with ice-cold assay buffer twice. The cells were removed from dishes by scratching and rinsing with 300 μl assay buffer. The cell suspension was added into vial with scintillator for liquid scintillation counting.

## Acknowledgments

The authors thank Drs. Ming Zhou, Ming Lei, Chaojun Li and Lijun Wang (for scientific discussion); Rijing Liao (assistance with the data analysis). Funding: This work is supported by National Key Research and Development Program of China (2017YFC1001303 and 2018YFC1004704), National Natural Science Foundation of China (U1632132, 31670849 and 91853206), SHIPM-pi fund No. JY201804 and SHIPM-sigma fund No. 2018JC002 from Shanghai Institute of Precision Medicine, Ninth People`s Hospital Shanghai Jiao Tong University School of Medicine. This work is also supported by Innovative Research Team of High-level Local Universities in Shanghai. Author contributions: Y. C. and Li Z. initiated the study. Y. C., A. Q., J. Z., Lu. Z. and D. L. designed research and wrote the paper. Li Z. and Q. W. performed the purification and crystallization work. Li. Z., F. Z., and Q. Z. collected and analyzed the data. D. Y. and Y. C. determined the structure. Li Z. and Lu. Z. performed functional assay. Competing interests: Authors declare no competing interests. Data and materials availability: Shanghai Synchrotron Radiation Facility (SSRF) beamlines BL18U1 and BL19U1 are used for x-ray crystallography data collection. The coordinates are deposited at the PDB accession code: 6KXW.

## Author Contributions

Y. C. and Li Z. initiated the study. Y. C., A. Q., J. Z., Lu. Z. and D. L. designed research and wrote the paper. Li Z. and Q. W. performed the purification and crystallization work. Li. Z., F. Z., and Q. Z. collected and analyzed the data. D. Y. and Y. C. determined the structure. Li Z. and Lu. Z. performed functional assay.

## Supplementary Information

**Fig. S1.**
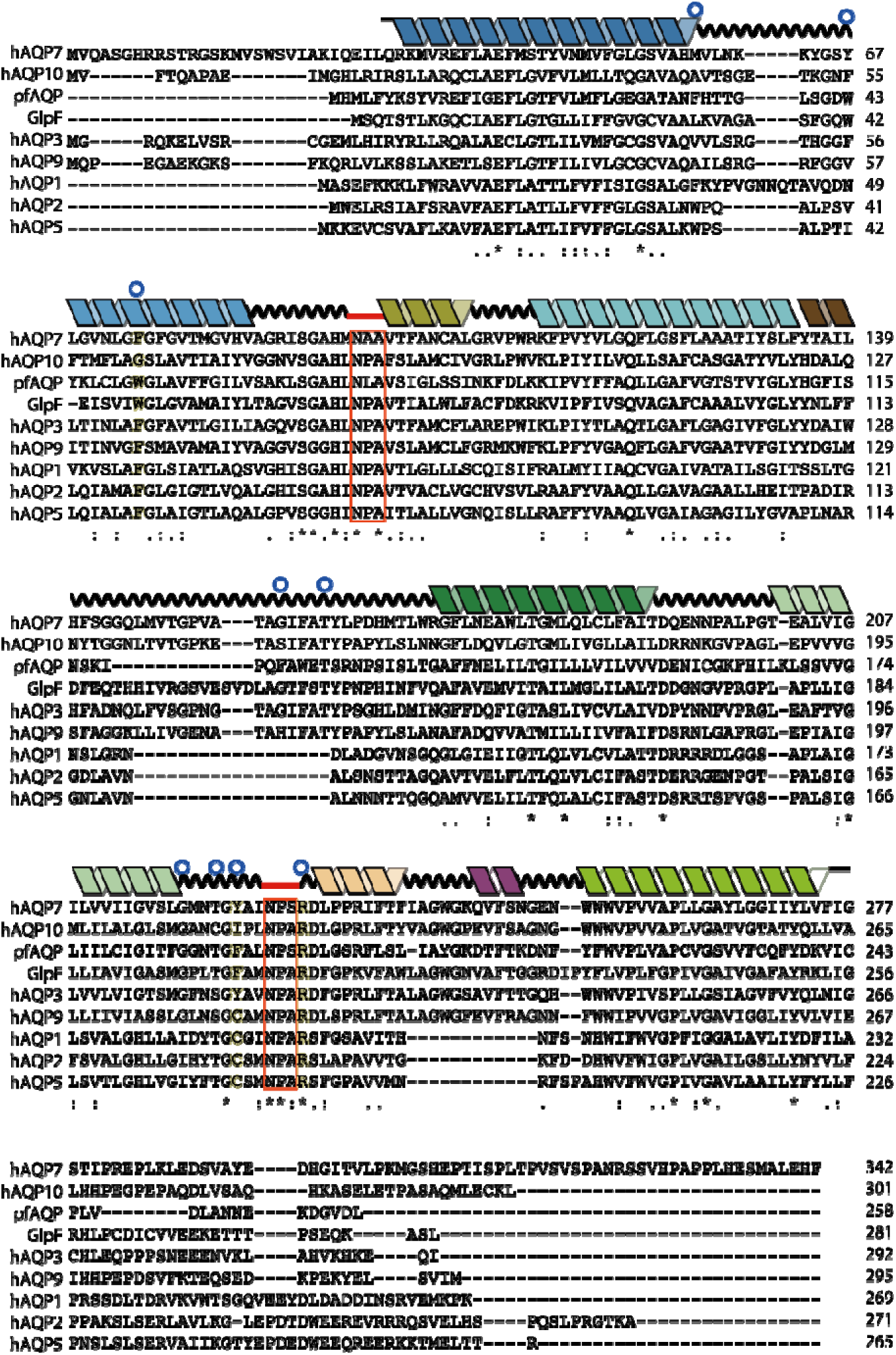
Amino acid sequence alignment and secondary structure of AQPs. Amino acid sequence alignment of Homo sapiens AQP7 (Uniprot ID O14520) with *Homo sapiens* AQP1 (Uniprot ID P29972), *Homo sapiens* AQP2 (Uniprot ID P41181), *Homo sapiens* AQP3 (Uniprot ID Q92482), *Homo sapiens* AQP5 (Uniprot ID P55064), *Homo sapiens* AQP9 (Uniprot ID O43315), *Escherichia coli* GlpF (Uniprot ID P0AER0), and *Plasmodium falciparum* AQP (Uniprot ID Q8WPZ6). AQP7 α-helices are shown as ribbons and colored according to the same scheme as in Fig 2A, with the loops shown as wave lines. The sequence similarity is shown with asterisks (*) for identical residues and colon (:) for strictly conserved residues. The residues forming Ar/R filter are marked with yellow background in sequence alignment and the “NPA” motifs marked with red frames. The residues in close proximity of glycerol molecule bound in AQP7 were labeled with blue circles.

**Fig. S2.**
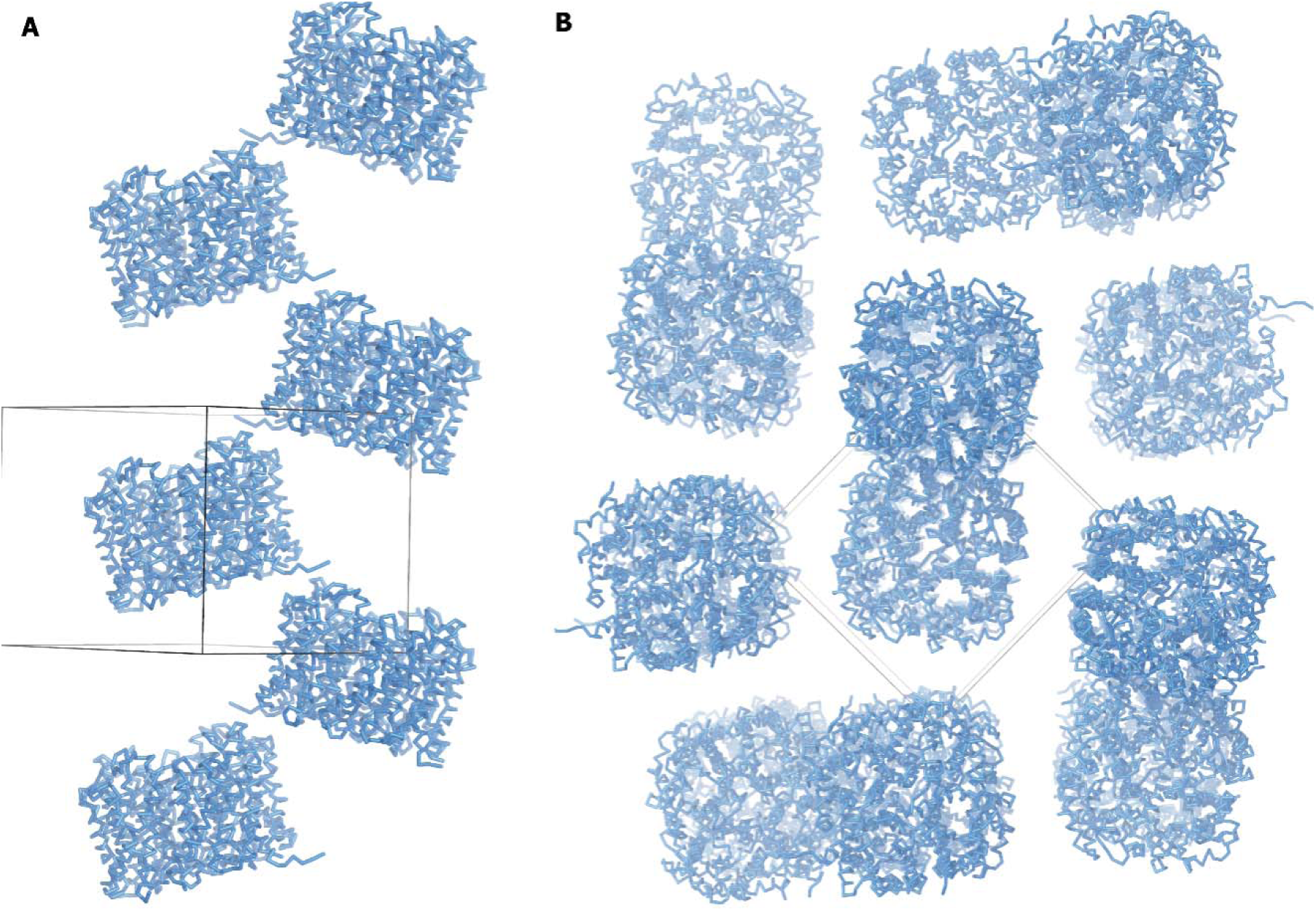
Crystal packing of AQP7. (A) crystal packing viewed from the side of the tetrameric AQP7. (A) crystal packing viewed from the top of the tetrameric AQP7.

**Fig. S3.**
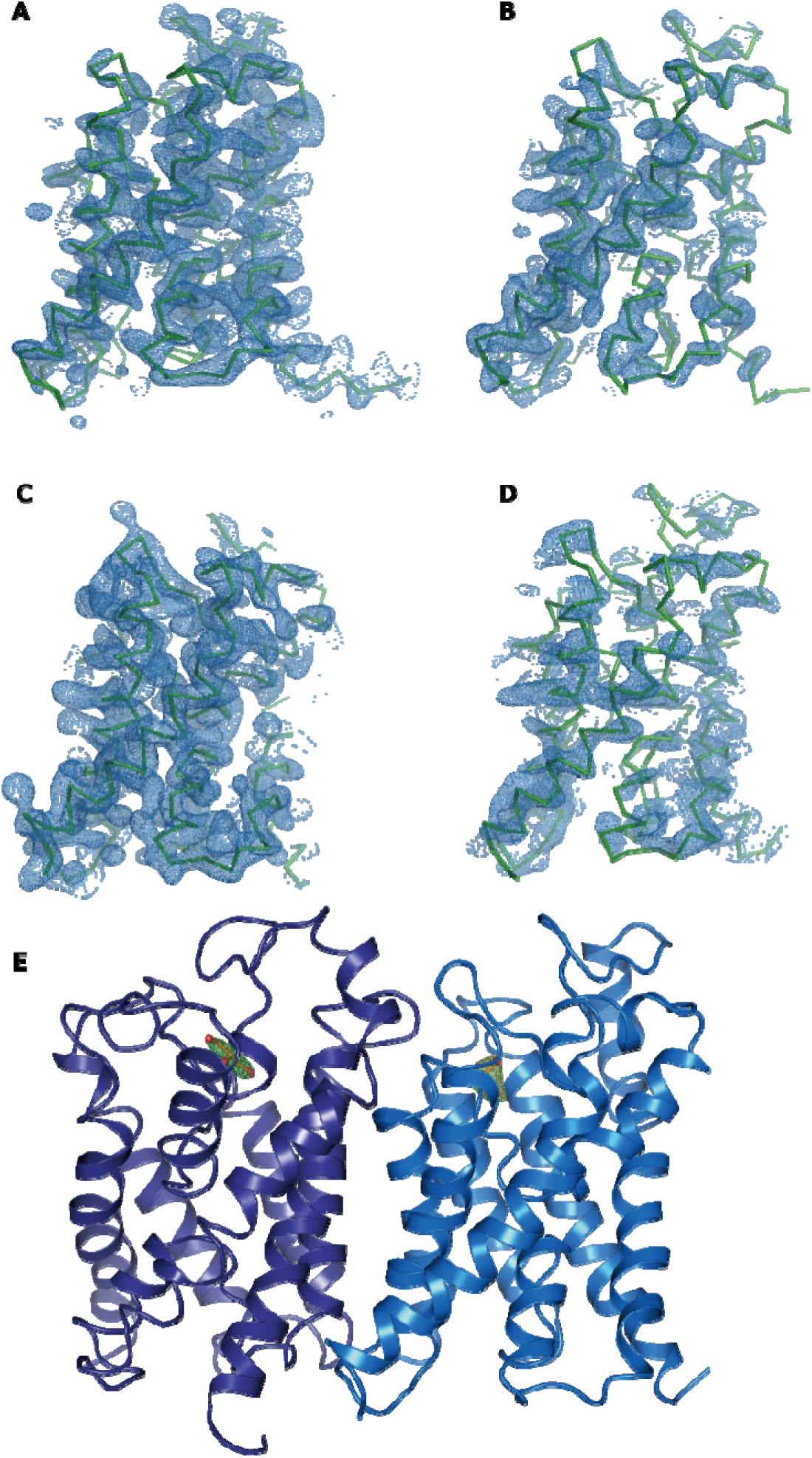
The comparison on electron density for protomers in AQP7 homotetramer. (A)-(D) the models and the corresponding electron density for chain A-C of AQP7 crystal structure. The Fo-Fc electron density map was contoured at 2.0 σ for chain A and C, 1.8 σ for chain B and D. (E) The electron density of glycerol bound in chain A and C. The AQP7 protein molecules were shown as cartoon representation in purple and blue, respectively and the chain B and D were removed for a clear observation.

**Table S1.**
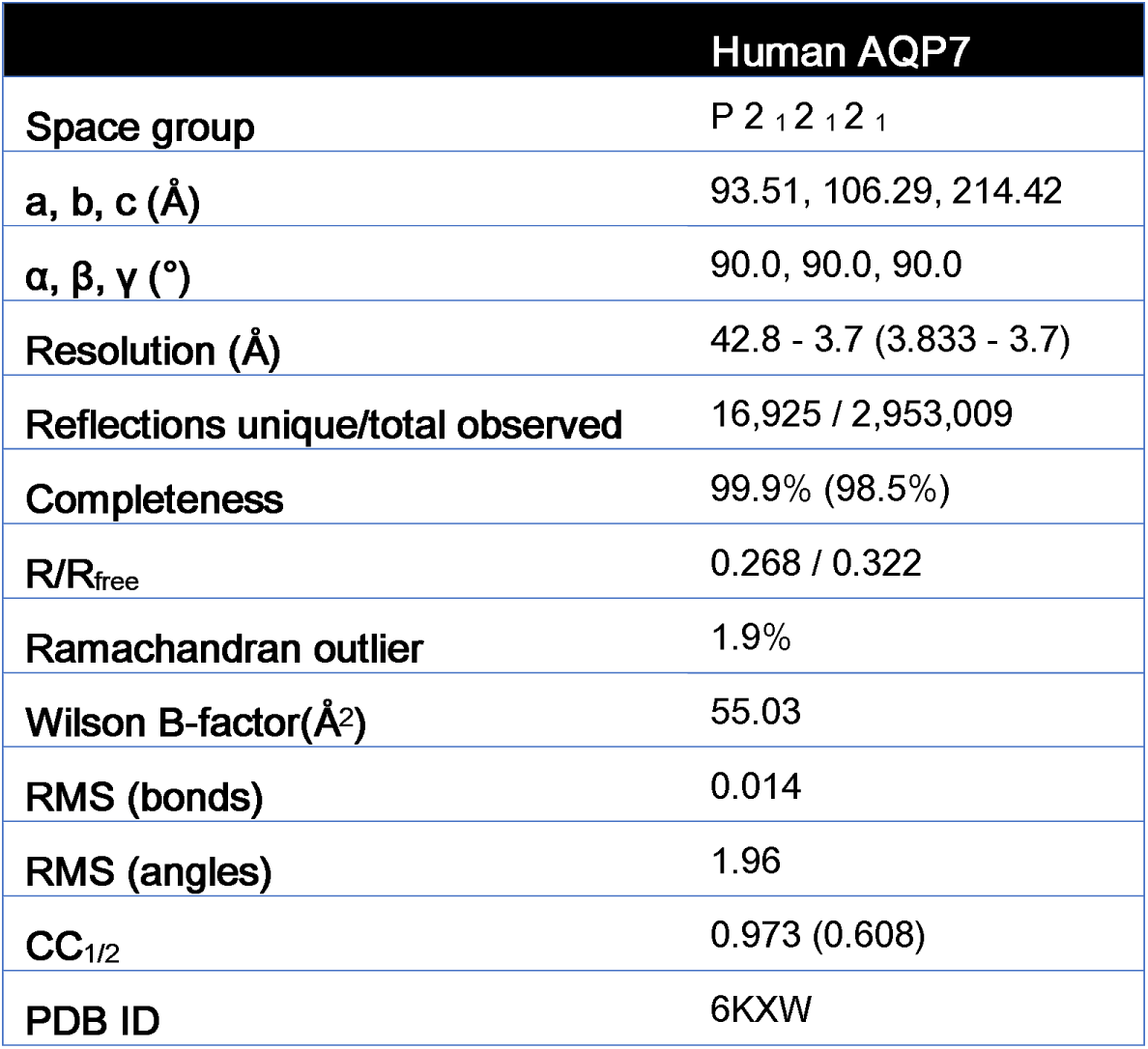

